## Supplementary material for "Generalizing the isothermal efficiency by using Gaussian distributions": Delila instructions for LexA sites

title 'LexA version = 3.23 of lexa.inst 2022 Aug 01'; (*
         2022 Aug 01, 3.23: clean up
         2022 Aug 01, 3.22: document delila
         2022 Aug 01, 3.21: add bibliography at end!!!
         2022 Aug 01, 3.20: Switch to version 3: NC_000913.3
                            I used sebo.p and seboxfr
         2022 Jul 31, 3.19: make instructions uniform for upgrade
         2020 Sep 11, 3.18: FernandezDeHenestrosa.Woodgate2000
         2011 Jun 03, 3.17: no version number on pieces from dbbk
         2011 Jun 02, 3.16: To track GenBank, rename:
                            E.coli-str.-K12-substr.-MG1655
                            to
                            E.coli-str.-K-12-substr.-MG1655
                            and add version number to piece.
         2008 Oct 20, 3.15: yebG
         2008 Oct 17, 3.14: official version
         2008 Oct 16, 3.13: switch to E.coli-str.-K12-substr.-MG1655
         2008 Apr 24, 3.12: dinD
         2008 Apr 24, 3.11: ydjM
         2008 Apr 23, 3.10: checking
         2008 Apr 19, 3.09: a bunch more ...
         2008 Apr 19, 3.08: finish recN
         2008 Apr 19, 3.07: umuD cleanup.
         2008 Apr 19, 3.06: umuD
         2008 Apr 18, 3.05: uvrD
         2008 Apr 17, 3.04: uvrD is next
         2008 Apr 17, 3.03: uvrA done, uvrB, sulA=sfiA
         2008 Apr 17, 3.02: uvrA started
         2008 Apr 17, 3.01: lexA and recA done
         2008 Apr 16, 3.00: rebuild entirely for E. coli genome.
         2006 Feb 20, 1.00: add direction to instructions
originally:
module lexa version = 1.21 of sitein 1991 Feb 5 *)

(* This file contains Delila instructions for grabbing the binding
sites of the LexA protein of E. coli.

The file is stored at:
https://alum.mit.edu/www/toms/lexa.inst

Information about how to use Delila is at:
https://alum.mit.edu/www/toms

References at the end are in Bibtex format for use with LaTeX.
https://alum.mit.edu/www/toms/latex.html

Thomas D. Schneider, Ph.D.
*)

set out-of-range reduce-range;

organism   E.coli-str.-K-12-substr.-MG1655;
chromosome E.coli-str.-K-12-substr.-MG1655;
piece NC_000913.3; (* complete genome *)

(****************************************************************)
(****************************************************************)
(* Proven LexA binding sites - DNase I or mutation **************)
(****************************************************************)
(****************************************************************)

(* lexA \cite{Little.Yanisch-Perron1981} figure 3B DNase I;
\cite{Brent.Ptashne1981} Figure 1 DNase I *)
(* original name: ecolexa *)
name "lexA-1"; get from 4257077 -200 to same +200 direction +;
name "lexA-1"; get from 4257078 +200 to same -200 direction -;
name "lexA-2"; get from 4257098 -200 to same +200 direction +;
name "lexA-2"; get from 4257099 +200 to same -200 direction -;

(* lexA \cite{Little.Yanisch-Perron1981} figure 3A DNase I;
\cite{Brent.Ptashne1981} Figure 1 DNase I *)
(* original name: ecoreca *)
name "recA"; get from 2823838 -200 to same +200 direction +;
name "recA"; get from 2823839 +200 to same -200 direction -;

(* uvrA \cite{Sancar.Mount1982} figure 1 DNase I
ssb is in the opposite direction.
*)

(* original name: ecouvra DNase I *)
name "uvrA"; get from 4273964 -200 to same +200 direction +;
name "uvrA"; get from 4273965 +200 to same -200 direction -;

(* uvrB \cite{Sancar.Rupp1982} figure 2 DNase I *)
(* original name: ecouvrb *)
name "uvrB"; get from 813441 -200 to same +200 direction +;
name "uvrB"; get from 813442 +200 to same -200 direction -;

(* ompA *)
(* This site is right in front of sulA.  ompA is downstream of sulA
(1020142).  So sulA is now listed by everyone but not ompA (1019276). 
There is no lexA site >10 bits between the end of sulA and the start
of ompA.  There is only a bad (black letter) 3 bit lexA site between
them.  So this was misnamed when I first did it! *)
(* original name: ecoompaii *)
{NAME "ompA"; GET FROM 1019276 -200 to same +200 direction +;}
{NAME "ompA"; GET FROM 1019277 +200 to same -200 direction -;}
(* I capitalized the instructions so they will never be recognized. *)

(* sulA Cole1983 Figure 4b \cite{Mizusawa.Gottesman1983} consensus match on
known promoter.  This was originally called ecoompaii, see above. 
Original name: sulA Berg1988e.  So there was a duplication in my
previous list!! tactgtacatccatacagta were identical.
Also sulA = sfiA (synonym) *)
name "sulA"; get from 1020951 -200 to same +200 direction +;
name "sulA"; get from 1020952 +200 to same -200 direction -;

(* uvrD \cite{Easton.Kushner1983} figure 5 DNase I *)
(* original name: uvrD Berg1988e *)
name "uvrD"; get from 3997916 -200 to same +200 direction +;
name "uvrD"; get from 3997917 +200 to same -200 direction -;

(* umuD \cite{Kitagawa.Kato1985} figure 6 DNase I *)
(* original name: umuDC Berg1988e *)
name "umuD"; get from 1230737 -200 to same +200 direction +;
name "umuD"; get from 1230738 +200 to same -200 direction -;

(* recN \cite{Rostas.Lloyd1987} figure 2 DNase I but data not shown
for recN-1 and recN-2 (see page 5048) and EMSA figure 3.  Though
Rostas.Lloyd1987 don't seem to notice, there are three discrete bands
in the EMSA.  So there clearly are three sites.  No other data
are available for recN-3, it is mentioned in
\cite{FernandezDeHenestrosa.Woodgate2000}.
RecN-3 is listed in the database of \cite{Robison.Church1998}
http://arep.med.harvard.edu/dpinteract/ but the references do not work
anymore.
---
\cite{Lewis.Mount1994} (page 510) point to
\cite{Schnarr.Granger-Schnarr1991} who say it is recN-3 is
cooperatively bound (page 427).  They say "M Kazmaier, MGS and MS in
preparation", however PubMed search for "Kazmaier Schnarr" on 2008 Apr
23 gave no additional papers! *)
(* original name: recN-1 Berg1988e *)
(* original name: recN-2 Berg1988e *)
name "recN-1"; get from 2751736 -200 to same +200 direction +;
name "recN-1"; get from 2751737 +200 to same -200 direction -;
name "recN-2"; get from 2751758 -200 to same +200 direction +;
name "recN-2"; get from 2751759 +200 to same -200 direction -;
name "recN-3"; get from 2751776 -200 to same +200 direction +;
name "recN-3"; get from 2751777 +200 to same -200 direction -;

(* ruv \cite{Shinagawa.Nakata1988} figure 7 DNase I, figure 2 sequence
*)
name "ruv"; get from 1946035 -200 to same +200 direction +;
name "ruv"; get from 1946036 +200 to same -200 direction -;

(* sbmC \cite{Baquero.Moreno1995} mutation.
See also \cite{Courcelle.Hanawalt2001} Table 1 *)
name "sbmC"; get from 2081286 -200 to same +200 direction +;
name "sbmC"; get from 2081287 +200 to same -200 direction -;

(* polB \cite{Lewis.Mount1994} Figure 1a EMSA, Table 2
polB = dinA *)
name "polB=dinA"; get from 65843 -200 to same +200 direction +;
name "polB=dinA"; get from 65844 +200 to same -200 direction -;

(* ftsK \cite{Lewis.Mount1994} Figure 1b EMSA, Table 2
ftsK = dinH.  See \cite{Lewis.Mount1994} - dinH is downstream
of lrp and that gene is now called ftsK.  *)
name "ftsK=dinH"; get from 933137 -200 to same +200 direction +;
name "ftsK=dinH"; get from 933138 +200 to same -200 direction -;

(* dinI \cite{Lewis.Mount1994} Figure 3 EMSA, Figure 4 EMSA, Table 2
dinI is downstream of pyrC, see \cite{Yasuda.Ohmori1996} *)
name "dinI"; get from 1121516 -200 to same +200 direction +;
name "dinI"; get from 1121517 +200 to same -200 direction -;

(* sosC \cite{Lewis.Mount1994} Figure 7a EMSA, Figure 9 *)
name "sosC=hsdS"; get from 4579925 -200 to same +200 direction +;
name "sosC=hsdS"; get from 4579926 +200 to same -200 direction -;

(* sosD \cite{Lewis.Mount1994} Figure 7b EMSA, Figure 9 *)
name "sosD"; get from 1823491 -200 to same +200 direction +;
name "sosD"; get from 1823492 +200 to same -200 direction -;

(* dinG \cite{Lewis.Mount1994} Table 2;
\cite{Lewis.Mount1992b} Table 1 mutations; Figure 1 EMSA.  *)
name "dinG"; get from 833045 -200 to same +200 direction +;
name "dinG"; get from 833046 +200 to same -200 direction -;

(* ydjM \cite{Fernandez_De_Henestrosa.Woodgate2000} EMSA: Figure 4,
   DNase I: Figure 5. *)
name "ydjM-1"; get from 1810186 -200 to same +200 direction +;
name "ydjM-1"; get from 1810187 +200 to same -200 direction -;
name "ydjM-2"; get from 1810168 -200 to same +200 direction +;
name "ydjM-2"; get from 1810169 +200 to same -200 direction -;

(*
@ NC_000913 1808192.0 +1 "lexA" "10.8 bits @ 1808192" 10.837679 -2.701801 0.003448
@ NC_000913 1808210.0 +1 "lexA" "17.8 bits @ 1808210" 17.841963 -0.876828 0.190290
@ NC_000913 1808418.0 +1 "lexA" " 8.7 bits @ 1808418" 8.674640 -3.265383 0.000547
*)

(************************************************************************)
(************************************************************************)
(* Genes known to be under LexA control with moderate experimental data *)
(************************************************************************)
(************************************************************************)

(* dinD \cite{Yasuda.Ohmori1996} \cite{Ohmori.Nagai1995}
pcsA = dinD.  Listed in the database of \cite{Robison.Church1998} and
listed in Table 1 of \cite{Fernandez_De_Henestrosa.Woodgate2000}
without a reference!  Lewis.Mount1994 Figure 9 and Figure 10.  This is
supposed to be a real site, but there is no direct evidence!  However
the indirect evidence is quite strong.  Ohmori.Nagai1995 (page 160)
measured control with fused lacZ, 3816132 is the location of the AluI
site.  The region 3815508 to 3816137 covers AluI region that was
induced.  There really is only one strong site in the region, about 24
bits.  There is also a weaker 6 bit site upstream.
*)
name "dinD"; get from 3817706 -200 to same +200 direction +;
name "dinD"; get from 3817707 +200 to same -200 direction -;

(* yebG
\cite{Lomba.de_Almeida1997} (59 bp sequence)
\cite{Kuhner.Gaub2004} (AFM data on sequence that is too short).
*)
name "yebG"; get from 1930774 -200 to same +200 direction +;
name "yebG"; get from 1930775 +200 to same -200 direction -;

(**********************************************************************)
(**********************************************************************)
(* Genes known to be under LexA control but no experimental data ******)
(**********************************************************************)
(**********************************************************************)

(* uvrC nocite{Stark.Moses1989} Figure 2, Figure3 EMSA.
nocite{Sharma.Moses1986} sequence data.
No LexA model fell into the right places.
The data are unclear.  DO NOT USE.
   NAME "uvrC"; GET FROM 1994100 -200 to same +200 direction +;
   NAME "uvrC"; GET FROM 1994100 +200 to same -200 direction -;
I capitalized the instructions so they will never be recognized.
*)

(***********************************************************************)
(***********************************************************************)
(* Sites dropped because they are not on the E. coli Chromosome ********)
(***********************************************************************)
(***********************************************************************)

(* mucAB - Plasmid
J Bacteriol. 1990 Nov;172(11):6223-31.
LexA-independent expression of a mutant mucAB operon.
McNally KP, Freitag NE, Walker GC.
"pKM101 is a naturally occurring plasmid that carries mucAB, an analog
of the umuDC operon, ..."
*)
(* original name: mucAB Berg1988e *)  (* DROP: Plasmid *)
(* original name: cle1-1 Berg1988e *) (* DROP: Plasmid *)
(* original name: cle1-2 Berg1988e *) (* DROP: Plasmid *)
(* original name: Col1b Berg1988e *)  (* DROP: Plasmid *)
(* original name: ColA-1 Berg1988e *) (* DROP: Plasmid *)
(* original name: ColA-2 Berg1988e *) (* DROP: Plasmid *)
(* original name: ColE2 Berg1988e *)  (* DROP: Plasmid *)
(* original name: clodf13pro  *)      (* DROP: Plasmid *)

(****************************************************************)
(****************************************************************)

(* other sites *)

(* \cite{Lewis.Mount1994} Table 2 (no sites)
Mutat Res. 1989 Nov;218(3):207-10.  The LexA protein does not bind
specifically to the two SOS box-like sequences immediately 5' to the
phr gene.  Payne NS, Sancar A.
NAME "phr"; GET FROM 740149 -200 to same +200 direction +; *)
(* I capitalized the instruction so it will never be recognized. *)

(* no sites found by model lexA 1.00 within 500 bp of start name
"recQ"; GET FROM 740149 -200 to same +200 direction +; See:
Lewis.Mount1994 page 510: "the non-canonical site near recQ ... has
not been shown to bind LexA" *)
(* I capitalized the instruction so it will never be recognized. *)

(*

% title 'LexA version = 3.20 of lexa.inst 2022 Aug 01';

@article{aaaaa,
author = "A. A. Aaaaa",
title = "{version = 56.98 of all.bib 2022 Jul 30}",
journal = "Tom Schneider's all.bib BibTeX reference Database",
comment = "MISSING object \rule{0.5em}{1ex}",
anothercomment = "the year is set to 1900 to provide a lower endpoint for
the year program",
year = "1900"}
--

@article{Baquero.Moreno1995,
author = "M. R. Baquero
 and M. Bouzon
 and J. Varea
 and F. Moreno",
title = "{\emph{sbmC}, a stationary-phase induced SOS
\emph{Escherichia coli} gene, whose product protects cells from the
DNA replication inhibitor microcin B17}",
journal = "Mol. Microbiol.",
volume = "18",
pages = "301--311",
pmid = "8709849",
year = "1995"}

@article{Brent.Ptashne1981,
author = "R. Brent
 and M. Ptashne",
title = "Mechanism of action of the {{\em lexA}} gene product",
journal = "Proc. Natl. Acad. Sci. USA",
volume = "78",
pages = "4204--4208",
comment = "original name: Brent1981",
year = "1981"}

@article{Courcelle.Hanawalt2001,
author = "J. Courcelle
 and A. Khodursky
 and B. Peter
 and P. O. Brown
 and P. C. Hanawalt",
title = "{Comparative gene expression profiles following UV exposure
in wild-type and SOS-deficient \emph{Escherichia coli}}",
journal = "Genetics",
volume = "158",
pages = "41--64",
pmid = "11333217",
pmcid = "PMC1461638",
year = "2001"}

@article{Easton.Kushner1983,
author = "A. M. Easton
 and S. R. Kushner",
title = "{Transcription of the \emph{uvrD} gene of \emph{Escherichia
coli} is controlled by the \emph{lexA} repressor and by attenuation}",
journal = "Nucleic Acids Res.",
volume = "11",
pages = "8625--8640",
pmid = "6324092",
pmcid = "PMC326612",
year = "1983"}

@article{FernandezDeHenestrosa.Woodgate2000,
author = "A. R. {Fernandez De Henestrosa}
 and T. Ogi
 and S. Aoyagi
 and D. Chafin
 and J. J. Hayes
 and H. Ohmori
 and R. Woodgate",
title = "{Identification of additional genes belonging to the LexA
regulon in \emph{Escherichia coli}}",
journal = "Mol. Microbiol.",
volume = "35",
pages = "1560--1572",
pmid = "10760155",
year = "2000"}

Fernandez_De_Henestrosa.Woodgate2000 not found in /Users/schneidt/papers/references/all.bib

@article{Kitagawa.Kato1985,
author = "Y. Kitagawa
 and E. Akaboshi
 and H. Shinagawa
 and T. Horii
 and H. Ogawa
 and T. Kato",
title = "{Structural analysis of the \emph{umu} operon required for
inducible mutagenesis in \emph{Escherichia coli}}",
journal = "Proc. Natl. Acad. Sci. USA",
volume = "82",
pages = "4336--4340",
pmid = "2989817",
pmcid = "PMC390408",
note = "\url
{https://doi.org/10.1073/pnas.82.13.4336}",
comment = "2021/06/06 21:55:47 formerly Kitagawa1985",
year = "1985"}

@article{Kuhner.Gaub2004,
author = "F. {K\"{u}hner}
 and L. T. Costa
 and P. M. Bisch
 and S. Thalhammer
 and W. M. Heckl
 and H. E. Gaub",
title = "{LexA-DNA bond strength by single molecule force spectroscopy}",
journal = "Biophys J",
volume = "87",
pages = "2683--2690",
pmid = "15454462",
pmcid = "PMC1304687",
year = "2004"}

@article{Lewis.Mount1992b,
author = "L. K. Lewis
 and D. W. Mount",
title = "{Interaction of LexA repressor with the asymmetric
\emph{dinG} operator and complete nucleotide sequence of the gene}",
journal = "J. Bacteriol.",
volume = "174",
pages = "5110--5116",
pmid = "1629168",
pmcid = "PMC206328",
year = "1992"}

@article{Lewis.Mount1994,
author = "L. K. Lewis
 and G. R. Harlow
 and L. A. Gregg-Jolly
 and D. W. Mount",
title = "{Identification of high affinity binding sites for LexA which
define new DNA damage-inducible genes in \emph{Escherichia coli}}",
journal = "J. Mol. Biol.",
volume = "241",
pages = "507--523",
pmid = "8057377",
note = "\url
{https://doi.org/10.1006/jmbi.1994.1528}",
comment = "2008Apr17_18:34:420 from pdf date",
year = "1994"}

@article{Little.Yanisch-Perron1981,
author = "J. W. Little
 and D. W. Mount
 and C. R. Yanisch-Perron",
title = "{Purified \emph{lexA} protein is a repressor of the
\emph{recA} and \emph{lexA} genes}",
journal = "Proc. Natl. Acad. Sci. USA",
volume = "78",
pages = "4199--4203",
pmid = "7027255",
pmcid = "PMC319756",
year = "1981"}

@article{Lomba.de_Almeida1997,
author = "M. R. Lomba
 and A. T. Vasconcelos
 and A. B. Pacheco
 and D. F. {de Almeida}",
title = "{Identification of \emph{yebG} as a DNA damage-inducible
\emph{Escherichia coli} gene}",
journal = "FEMS Microbiol. Lett.",
volume = "156",
pages = "119--122",
pmid = "9368369",
year = "1997"}

@article{Mizusawa.Gottesman1983,
author = "S. Mizusawa
 and D. Court
 and S. Gottesman",
title = "{Transcription of the \emph{sulA} gene and repression by LexA}",
journal = "J. Mol. Biol.",
volume = "171",
pages = "337--343",
pmid = "6317868",
year = "1983"}

@article{Ohmori.Nagai1995,
author = "H. Ohmori
 and M. Saito
 and T. Yasuda
 and T. Nagata
 and T. Fujii
 and M. Wachi
 and K. Nagai",
title = "{The \emph{pcsA} gene is identical to \emph{dinD} in
\emph{Escherichia coli}}",
journal = "J. Bacteriol.",
volume = "177",
pages = "156--165",
pmid = "8002613",
pmcid = "PMC176568",
year = "1995"}

@article{Robison.Church1998,
author = "K. Robison
 and A. M. McGuire
 and G. M. Church",
title = "{A comprehensive library of DNA-binding site matrices for 55
proteins applied to the complete \emph{Escherichia coli} K-12
genome}",
journal = "J. Mol. Biol.",
volume = "284",
pages = "241--254",
pmid = "9813115",
note = "http://arep.med.harvard.edu/dpinteract/",
year = "1998"}

@article{Rostas.Lloyd1987,
author = "K. Rostas
 and S. J. Morton
 and S. M. Picksley
 and R. G. Lloyd",
title = "{Nucleotide sequence and LexA regulation of the
\emph{Escherichia coli} \emph{recN} gene}",
journal = "Nucleic Acids Res.",
volume = "15",
pages = "5041--5049",
pmid = "3037486",
pmcid = "PMC305946",
year = "1987"}

@article{Sancar.Mount1982,
author = "A. Sancar
 and G. B. Sancar
 and W. D. Rupp
 and J. W. Little
 and D. W. Mount",
title = "{LexA protein inhibits transcription of the \emph{E. coli}
\emph{uvrA} gene \emph{in vitro}}",
journal = "Nature",
volume = "298",
pages = "96--98",
pmid = "6283374",
year = "1982"}

@article{Sancar.Rupp1982,
author = "G. B. Sancar
 and A. Sancar
 and J. W. Little
 and W. D. Rupp",
title = "{The \emph{uvrB} gene of \emph{Escherichia coli} has both
\emph{lexA}-repressed and \emph{lexA}-independent promoters}",
journal = "Cell",
volume = "28",
pages = "523--530",
pmid = "6280873",
year = "1982"}

@article{Schnarr.Granger-Schnarr1991,
author = "M. Schnarr
 and P. Oertel-Buchheit
 and M. Kazmaier
 and M. Granger-Schnarr",
title = "{DNA binding properties of the LexA repressor}",
journal = "Biochimie",
volume = "73",
pages = "423--431",
pmid = "1911942",
year = "1991"}

@article{Sharma.Moses1986,
author = "S. Sharma
 and T. F. Stark
 and W. G. Beattie
 and R. E. Moses",
title = "{Multiple control elements for the \emph{uvrC} gene unit of
\emph{Escherichia coli}}",
journal = "Nucleic Acids Res.",
volume = "14",
pages = "2301--2318",
pmid = "3515318",
pmcid = "PMC339659",
year = "1986"}

@article{Shinagawa.Nakata1988,
author = "H. Shinagawa
 and K. Makino
 and M. Amemura
 and S. Kimura
 and H. Iwasaki
 and A. Nakata",
title = "{Structure and regulation of the \emph{Escherichia coli}
\emph{ruv} operon involved in DNA repair and recombination}",
journal = "J. Bacteriol.",
volume = "170",
pages = "4322--4329",
pmid = "2842314",
pmcid = "PMC211445",
year = "1988"}

@article{Stark.Moses1989,
author = "T. Stark
 and R. E. Moses",
title = "{Interaction of the LexA repressor and the \emph{uvrC}
regulatory region}",
journal = "FEBS Lett",
volume = "258",
pages = "39--41",
pmid = "2556297",
year = "1989"}

@article{Yasuda.Ohmori1996,
author = "T. Yasuda
 and T. Nagata
 and H. Ohmori",
title = "{Multicopy suppressors of the cold-sensitive phenotype of the
\emph{pcsA68} (\emph{dinD68}) mutation in \emph{Escherichia coli}}",
journal = "J. Bacteriol.",
volume = "178",
pages = "3854--3859",
pmid = "8682790",
pmcid = "PMC232646",
year = "1996"}

*)
